# Happens in the best of subfamilies: Establishment and repeated replacements of co-obligate secondary endosymbionts within Lachninae aphids

**DOI:** 10.1101/059816

**Authors:** Alejandro Manzano-Marín, Gitta Szabo, Jean-Christophe Simon, Matthias Horn, Amparo Latorre

**Affiliations:** Institut Cavanilles de Biodiversitat i Biologia Evolutiva, Universitat de València, Paterna, Comunitat Valenciana, Spain; Department of Microbiology and Ecosystem Science, University of Vienna, Vienna, Austria; UMR1349 Institut de Génétique, Environnement et Protection des Plantes (IGEPP), Institut National de la Recherche Agronomique (INRA), Le Rheu, Bretagne, France; Área de Genómica y Salud de la Fundación para el fomento de la Investigación Sanitaria y Biomédica de la Comunitat Valenciana (FISABIO)-Salud Pública, València, Comunitat Valenciana, Spain

**Keywords:** Serratia symbiotica, Lachninae, Candidatus Fukatsuia symbiotica, aphid endosymbiont, symbiont replacement

## Abstract

Virtually all aphids maintain an obligate mutualistic symbiosis with bacteria from the *Buchnera* genus, which produce essential nutrients for their aphid hosts. Most aphids from the Lachninae subfamily have been consistently found to house additional endosymbionts, mainly *Serratia symbiotica*. This apparent dependence on secondary endosymbionts was proposed to have been triggered by the loss of the riboflavin biosynthetic capability by *Buchnera* in the Lachninae last common ancestor. However, an integral large-scale analysis of secondary endosymbionts in the Lachninae is still missing, hampering the interpretation of the evolutionary and genomic analyses of these endosymbionts. Here, we analysed the endosymbionts of selected representatives from seven different Lachninae genera and nineteen species, spanning four tribes, both by FISH (exploring the symbionts’ morphology and tissue tropism) and 16S rRNA gene sequencing. We demonstrate that all analysed aphids possess dual symbiotic systems, and while most harbour *S. symbiotica*, some have undergone symbiont replacement by other phylogenetically-distinct bacterial taxa. We found that these secondary associates display contrasting cell shapes and tissue tropism, and some appear to be lineage-specific. a scenario for symbiont establishment in the Lachninae, followed by changes in the symbiont’s tissue tropism and symbiont replacement events, thereby highlighting the extraordinary versatility of host-symbiont interactions.

**Originality-Significance Statement:** A key question in evolutionary biology is that of how mutualism evolves. One way to approach this problem is to investigate recently-established mutualistic associations, particularly by comparing various symbiotic systems in closely related hosts. Here, we present a most comprehensive study to investigate co-obligate symbioses in aphids, focusing in the Lachninae subfamily. While most aphids keep an obligate vertically-transmitted association with intracellular *Buchnera* bacteria, some, such as members of the Lachninae subfamily, host an additional putative co-obligate symbiont. Thus, the Lachninae dual symbiotic systems offer a unique opportunity to understand the evolutionary dynamics of host-symbiont associations, in particularly how secondary symbionts become obligate and eventually may be replaced. Through genome sequencing of three aphid species belonging to distantly related tribes within the subfamily, we have previously corroborated that they have indeed established co-obligate mutualistic associations with the *S. symbiotica* secondary endosymbiotic bacterium. This was putatively facilitated by an ancient pseudogenisation of the riboflavin biosynthetic pathway in *Buchnera*, rendering it unable to provide the essential vitamin to the host. However, not all Lachninae members harbour *S. symbiotica*, some species being associated to at least four different bacterial taxa. To correctly interpret the genomic data and to understand the evolutionary dynamics of these symbiotic associations, a wide-range analysis of both the phylogenetic relations as well as of the secondary symbionts’ localisation within the bacteriome is needed. To tackle this, we have combined phylogenetic analyses of the symbionts’ 16S rRNA gene sequences and FISH microscopy, to understand the symbiont’s identity as well as the morphological characteristics and tissue tropism. The phylogenetic affinities and patterns of co-divergence of the symbionts, in combination with previously published genomic data, have enabled us to build an evolutionary scenario for the establishment, changes in tissue tropism such as “stable” internalisation into bacteriocytes, and replacements of the putative “ancient” secondary endosymbiont from the Lachninae last common ancestor. Also, we were able to determine through phylogenetic analyses that some putative co-obligate endosymbionts may have evolved from once facultative ones. The evolutionary framework presented here reveals a dynamic pattern for the more recent evolutionary history of these symbioses, including replacement and novel acquisition of phylogenetically different co-obligate symbionts. This study opens new research avenues on this symbiont-diverse subfamily, providing insight into how mutualism in endosymbiotic associations can evolve, and the role these bacteria have played in the species’ adaptation and even in the speciation process.

## Introduction

The increasing recognition that symbionts play an important role in the ecology and evolution of their hosts, as well as the rapid changes in the type and nature of these symbiotic associations, call for an evolutionary framework to understand these dynamics. Symbiotic relationships between aphids and their primary obligate bacterial endosymbiont *Buchnera aphidicola*, represent one of the most well studied cases of bacterial endosymbiosis within animals. The *Buchnera* symbiosis is found across all modern aphids (Aphididae Latreille, 1802 family) (Buchner, 1953), with the notable exception of members belonging to the monophyletic Cerataphidini tribe Baker, 1920, in which *Buchnera* has been replaced by an extracellular yeast-like symbiont (Buchner, 1953; Fukatsu and Ishikawa, 1992; Fukatsu *et al.*, 1994). *Buchnera* cells have a round and pleomorphic shape (Michalik *et al.*, 2014), and inhabit the cytoplasm of bacteriocytes (specialised cells evolved to house the endosymbiont), which make up a distinct organ-like structure called the bacteriome (Buchner, 1953; Fukatsu *et al.*, 1998). The onset of the aphid-*Buchnera* symbiosis dates back to at least 80-150 million years ago (hereafter **Mya**) (von Dohlen and Moran, 2000). *Buchnera*, as other “ancient” obligate endosymbionts, underwent a rapid genome erosion early in its evolutionary history with aphids, resulting in a high degree of synteny among distantly related *Buchnera* (Tamas *et al.*, 2002; van Ham *et al.*, 2003), and since then, lineages of both partners have been co-diverging. This has been evidenced through phylogenetic reconstructions using *Buchnera* DNA or amino acid sequences, which parallel their aphid hosts’ evolutionary relationships (Munson *et al.*, 1991; Jousselin *et al.*, 2009; Liu *et al.*, 2013). Besides *Buchnera*, aphids can also harbour secondary endosymbionts (in addition to the primary symbiont), these being of facultative or obligate nature in some lineages. Contrary to obligate symbionts, facultative ones are not required for the correct development, reproduction and survival of their host. Still, they can provide a benefit under certain environmental or ecological conditions (conditional mutualism) (reviewed in Oliver *et al.*, 2010, 2014). To date, various secondary facultative bacterial endosymbionts have been identified, primarily in the pea aphid *Acyrthosiphon pisum* (Aphidinae subfamily) (Fukatsu *et al.*, 2000, 2001; Sakurai *et al.*, 2005; Degnan, Leonardo, *et al.*, 2009; Degnan, Yu, *et al.*, 2009; Guay *et al.*, 2009; Tsuchida *et al.*, 2014). These secondary symbionts have a very different tissue tropism than *Buchnera,* as they can be present in separate bacteriocytes (called secondary bacteriocytes), co-infecting the primary endosymbiont’s bacteriocytes, located in sheath cells (at the periphery of the bacteriome and found closely associated to bacteriocytes), and/or free in the haemocoel (Fukatsu *et al.*, 2000; Sandström *et al.*, 2001; Moran *et al.*, 2005; Sakurai *et al.*, 2005; Michalik *et al.*, 2014). While many secondary symbionts are facultative for their hosts, some seem to have established co-obligate associations with their respective symbiotic partners. In this respect, the subfamily Lachninae Herrich-Schaeffer, 1854 of aphids is peculiar, in that all members analysed thus far by microscopy techniques have been found to house secondary endosymbionts (Buchner, 1953; Fukatsu and Ishikawa, 1998; Fukatsu *et al.*, 1998; Lamelas *et al.*, 2008; Pyka-Fościak and Szklarzewicz, 2008; Michalik *et al.*, 2014).

The Lachninae can be divided into 5 monophyletic tribes: i) **Lachnini** Herrich-Schaeffer, 1854, ii) **Stomaphidini** Mordvilko, 1914, iii) **Tramini** Herrich-Schaeffer, 1854, iv) **Tuberolachnini** Mordvilko, 1942, and v) **Eulachnini** Baker, 1920 (Chen, Favret, *et al.*, 2015) (**Fig 1**). The first four tribes comprise 112 known species organised into 13 genera, however, due to the lack of molecular data, the phylogenetic affiliation of three of these (*Neonippolachnus*, *Sinolachnus*, and *Eotrama*) remains unresolved, and thus are not included in the Figure. While the Lachnini, Tramini, and Tuberolachnini feed on angiosperms, the *Stomaphidini* feed on both angiosperm and gymnosperm trees (bark-trunk and root). The latter have evolved some of the longest mouthparts among aphids, particularly the trunk-feeding species (Blackman and Eastop, 1994). The Tramini are unique in that they solely feed on the roots of herbaceous plants, mostly composites (Blackman and Eastop, 2006). Finally, the Eulachnini, which exclusively feed on conifers, are classified into 4 genera: *Essigella*, *Eulachnus*, *Pseudessigella* (with no molecular data available), and *Cinara.* The latter is the largest genus within the Lachninae and has been traditionally taxonomically classified into three subgenera (*Cinara*, *Cupressobium*, and *Cedrobium*). However, recent extensive molecular work on members of the subgenus *Cinara* has found the subgenus *Cinara* (*Cinara*) polyphyletic, and thus has divided the genus into three major phylogenetic clades, termed simply **A**, **B**, and **C** (Meseguer *et al.*, 2015).

**Fig 1.**
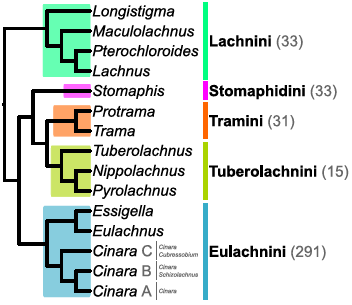
Dendrogram depicting the relationships among Lachninae aphids. Dendogram based on (Chen, Favret, *et al.*, 2015) and (Meseguer *et al.*, 2015). Coloured boxes shading, as well as vertical lines on the right-hand side, delimit the tribal clades. Names next to the coloured vertical lines provide tribal names and number of extant species (in brackets), according to (Favret, 2016). For *Cinara* clades, a grey vertical bar is followed by the different subgenera that make up the clades A, B, and C. The genera *Eotrama* (Tramini), *Neonippolachnus* (Tuberolachnini), *Synolachnus* (Tuberolachnini), and *Pseudessigella* (Eulachnini) are not included, as no molecular data for their reliable placement is currently available. Based on (Chen, Favret, *et al.*, 2015), in the current work we have considered the *Schizolachnus* genus as a subgenus within the *Cinara* clade B.

Known secondary symbionts of Lachninae differ in tissue tropism and cell shape (Buchner, 1953; Fukatsu and Ishikawa, 1998; Fukatsu *et al.*, 1998; Pyka-Fościak and Szklarzewicz, 2008; Michalik *et al.*, 2014), as well as in phylogenetic origin (Russell *et al.*, 2003; Lamelas *et al.*, 2008; Burke *et al.*, 2009; Jousselin *et al.*, 2016). Although different bacterial taxa have been found associated to Lachninae aphids, many species of this subfamily have been systematically found associated with members of the bacterial genus *Serratia*, mainly *Serratia symbiotica* (Fukatsu *et al.*, 1998; Russell *et al.* 2003; Lamelas *et al.*, 2008; Burke *et al.*, 2009; Chen *et al.* 2015; Jousselin *et al.*, 2016) (for a summary see **Table S1**). Particularly, most *Cinara* species have been consistently found to house *S. symbiotica* strains, which form two phylogenetically distinct clusters (based on 16S rRNA gene sequences), one “facultative-like” and one “obligate-like” (hereafter **FL** and **OL**, respectively) (Lamelas *et al.*, 2008; Burke *et al.*, 2009). Whole genome sequencing and metabolic reconstruction of the *Buchnera*-*S. symbiotica* bacterial consortia of three Lachninae species revealed that *S. symbiotica* strains, belonging to FL and OL, had indeed established co-obligate associations along with *Buchnera* in these hosts (Gosalbes *et al.*, 2008; Lamelas, Gosalbes, Manzano-Marín, *et al.*, 2011; Manzano-Marín and Latorre, 2014; Manzano-Marín *et al.*, 2016). Through comparative genomics of these di-(*Buchnera*-*S. symbiotica*) *vs.* mono-endosymbiotic (*Buchnera*-only) systems, it was postulated that the establishment of *S. symbiotica* as a co-obligate endosymbiont in the Lachninae last common ancestor (hereafter **LLCA**) was facilitated by a putative ancient pseudogenisation of the riboflavin biosynthetic pathway in *Buchnera* and the complementation of this loss-of-function by *S. symbiotica* (Manzano-Marín and Latorre, 2014; Manzano-Marín *et al.*, 2016). Substantial differences regarding cell morphology and genomic characteristics observed among extant *S. symbiotica* strains suggest that these represent different stages of the genome reduction process towards a highly reduced obligate intracellular symbiont.

Given the overwhelming evidence pointing towards a dependency on secondary endosymbionts within the Lachninae, we sought to further understand the evolutionary succession of establishments, replacements, and changes in tissue tropism (e.g. “stable” internalisations into distinct bacteriocytes), of secondary endosymbionts among species of this subfamily. For this purpose, we have identified the secondary endosymbionts of distantly related Lachninae aphids belonging to 19 different species (8 different genera collected in three different countries) (**Table S1**) through 16S rRNA gene sequencing and phylogenetic analysis. In selected species, we have determined the location of the secondary endosymbionts using fluorescence *in situ* hybridisation (**FISH**) with 16S rRNA targeted specific oligonucleotide probes. We propose an evolutionary scenario for the establishment of an original secondary co-obligate endosymbiont in the LLCA, followed by symbiont replacements, “stable” internalisations of these into distinct bacteriocytes, and/or the putative establishment of tertiary obligate symbionts in different aphid lineages from this symbiont-diverse subfamily.

## Results

### *S. symbiotica* and *S. marcescens*-like secondary symbionts

Most *Cinara* spp. investigated so far have been found to be associated with different *S. symbiotica* strains (Lamelas *et al.*, 2008; Burke *et al.*, 2009; Jousselin *et al.*, 2016). Here, we have collected 19 representatives (comprising 11 species) of *Cinara* clades A (n=5), B (n=7), and C (n=7), and have identified their endosymbionts through PCR, cloning, and sequencing of their 16S rRNA genes. We found that the secondary symbionts of all of the collected species – except for *Cinara* (*Cinara*) *confinis* (*Cinara* clade C) and *Cinara* (*Schizolachnus*) *obscurus* – were indeed affiliated with *S. symbiotica* (**Table S1**).

To test for co-speciation previously observed for *Buchnera*-*Serratia* symbiont pairs within the Lachninae in the light of our new data, we performed a Bayesian phylogenetic reconstruction using currently available 16S rRNA gene sequences of both *Buchnera* (**Fig 2A**) and *Serratia* (**Fig 2B**) from Lachninae aphids. Contrary to earlier studies (Lamelas *et al.*, 2008; Burke *et al.*, 2009), we failed to recover the previously described FL and OL *S. symbiotica* clusters. We found that all Lachninae *S. symbiotica* strains, but the one from *Trama caudata*, form a well-supported and unresolved monophyletic clade nested within a group composed mainly of facultative strains of *S. symbiotica* from Aphidinae aphids, a strain from *Adelges tsugae* (Hemiptera: Adelgidae), and one from the Lachninae aphid *Trama troglodytes* (**Fig 2B**). This unresolved clade contains the “early” co-obligate *S. symbiotica* strain from *Cinara (Cupressobium) tujafilina* (Manzano-Marín and Latorre, 2014) and a strain from the closely related *Cinara (Cupressobium) cupressi*. Within the Lachninae *S. symbiotica* clade, we recovered three well-supported monophyletic clades made up from: (i) *Cinara* (*Cinara*) ponderosae and *Cinara* (*Cinara*) *terminalis* (both from *Cinara* cluster A), (ii) some *Stomaphis* spp., and (iii) most *Lachnus* species. The latter belong to various closely related species, some suspected to be synonyms of *Lachnus tropicalis* (Blackman and Eastop, 1994), which would be consistent with the high sequence identity (>99%) of their *S. symbiotica* endosymbionts’ 16S rRNA gene. Interestingly, most *S. symbiotica* strains from *Cinara* clade A form a well-supported monophyletic clade, and within this, there is high congruency with the phylogenetic relationships of the *Buchnera* strains found in the respective hosts, particularly within two subclades (**Fig 2A** and **B**, vertical black lines). In contrast, most *S. symbiotica* strains from *Cinara* clade B are polyphyletic, and their phylogenetic relationships do not seem to mirror those of the respective *Buchnera* symbiont. Curiously, both the *Buchnera* and *S. symbiotica* from *Cinara* (*Cupressobium*) *costata* are recovered nested within strains from *Cinara* clade A. This contrasts with previously established *Cinara* phylogenetic relationships (Chen, Favret, *et al.*, 2015; Meseguer *et al.*, 2015), which show this species to be part of a basal group of *Cinara* clade C. On the other hand, *S. symbiotica* strains from *Lachnus roboris*, *Lachnus quercihabitans*, *Tuberolachnus salignus,* and *Pterochloroides persicae* are all recovered within a clade encompassing most *S. symbiotica* strains from *Cinara* spp., reflecting no congruency with neither their hosts’ nor their corresponding *Buchnera* relationships.

*Serratia* strains from *Stomaphis* spp. are recovered nested within both the free-living *S. marcescens* strains and the Lachninae *S. symbiotica* clade. The former constitutes what we denominate the ***S***. ***m****arcescens*-**l**ike **s**econdary **s**ymbionts (hereafter **SMLSS**), all of which have been identified from aphids belonging to a single clade of *Stomaphis* spp. (**Fig 2A** and **B**). The latter are recovered as a monophyletic clade which is congruent with the *Buchnera* phylogeny, and as basal to the clade comprising most *S. symbiotica* strains from *Cinara* species.

**Fig 2.**
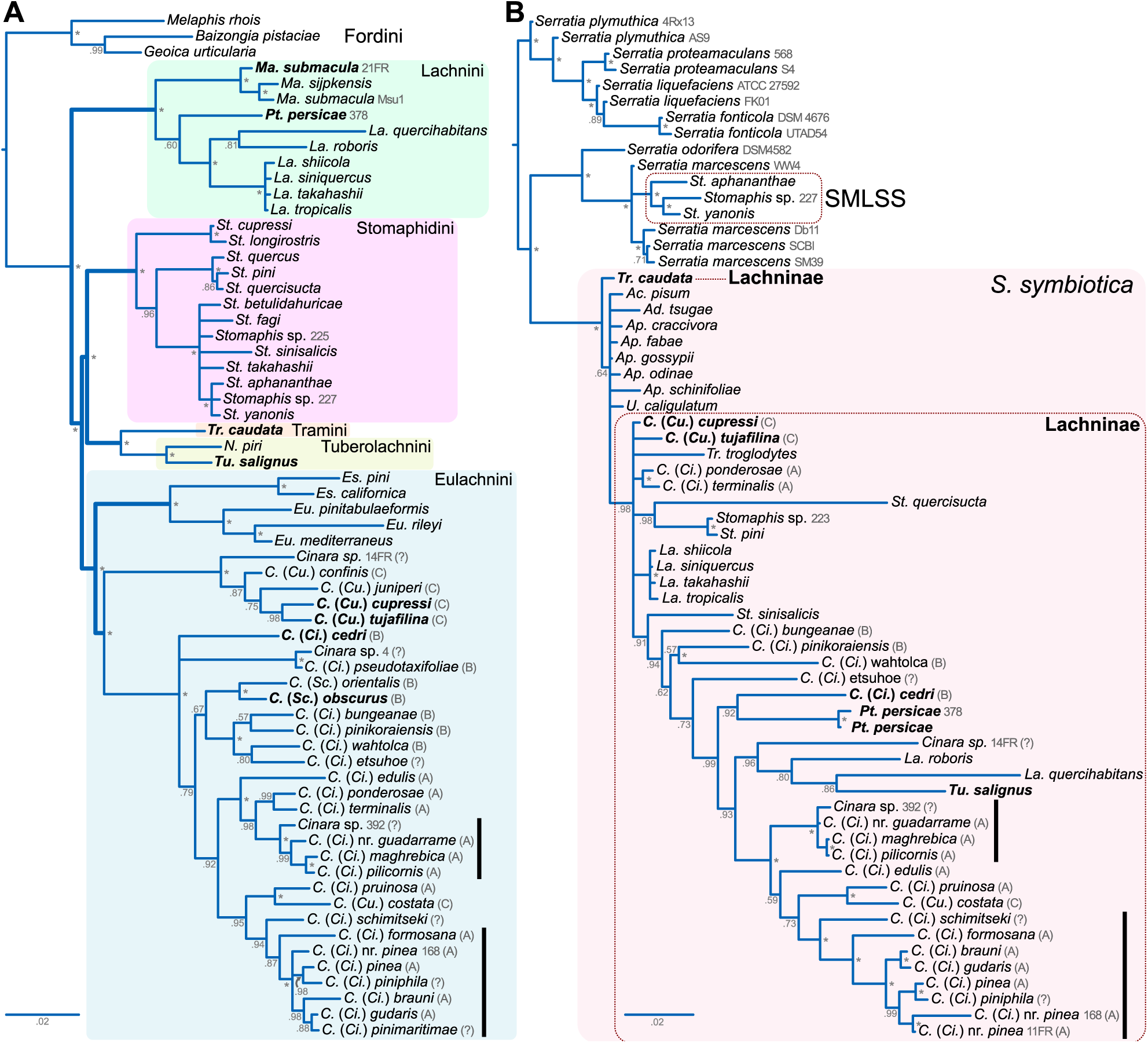
16S rRNA-based phylogenetic relationships of *Buchnera* and *Serratia* strains from Lachninae aphids. Bayesian phylogram of **(A)** *Buchnera* and **(B)** *Serratia* symbionts from selected aphids. *Buchnera* from the Fordini tribe and free-living *Serratia* strains were used for rooting the respective trees. Values at nodes indicate the posterior probability. An “*” at the node indicate a posterior probability of 1. For the *Buchnera* tree, the thicker branches represent constrained relationships within Lachninae tribes according to (Chen, Favret, *et al.*, 2015). Following the species name, strain/isolate names and corresponding *Cinara* clade are indicated in grey, without and with parenthesis, respectively. Aphid tribe names in **A** are indicated at the top-right of the coloured boxes. The coloured box in **B** delimits the *S. symbiotica* clade, while dotted boxes delimit the SMLSS (from *Stomaphis* spp.) and the *S. symbiotica* strains from Lachninae aphids, respectively. In both **A** and **B**, bold-lettered species names indicate the selected species we have used for FISH microscopy.

Next, we investigated the diversity in tissue tropism and intracellular location of *S. symbiotica* within distantly related Lachninae by whole-mount FISH using aphid embryos. We found that all of the FISH-analysed specimens for *S. symbiotica* indeed housed this symbiont, meaning they were fixed in the population and suggesting that they represent obligate symbionts. Interestingly, we observed a great diversity in both cell-shape and tissue tropism of *S. symbiotica* among the selected Lachninae (**Figs 3 and S1**). In *C.* (*Cu.*) *tujafilina* and *C.* (*Cu.*) *cupressi*, *S. symbiotica* is present in the periphery of the *Buchnera* bacteriocytes, co-infecting them, and also occupying its own bacteriocytes (**Figs 3A–B and S1A-D**). On the other hand, in *Cinara* (*Cinara*) *cedri* (clade B); *Tu. salignus; Pt. Persicae;* and *Tr. caudata*, *S. symbiotica* is housed exclusively inside distinct bacteriocytes (**Figs 3C, D, E, F, and S1E-L**), and thus “stably” internalised in its own distinct host cells. The distribution of these secondary bacteriocytes is different between the aphid species. In *C. (Ci.) cedri* they are interspersed among *Buchnera* bacteriocytes (**Figs 3C, and S1E-G**), in *Tu. salignus* they are found forming a “bacteriome core” surrounded by primary bacteriocytes (**Figs 3D and S1H-I**), and in *Pt. persicae* they form a “layer” along the bacteriome (**Figs 3E and S1J**). *S. symbiotica* cells appear round-shaped in *C. (Ci.) cedri*, *Tu. salignus*, and *Pt. Persicae*, while in *Tr. caudata* the secondary symbiont retains a rod shape and cell size similar to free-living *Serratia* strains (**Figs 3F and S1K-L**). Moreover, the *S. symbiotica* strains of *C.* (*Cu.*) *tujafilina* (**Figs 3A and S1A-C**) and *C.* (*Cu.*) *cupressi* (**Figs 3B and S1D**) show an elongated filamentous cell shape, similarly to the facultative *S. symbiotica* symbiont of *Ac. pisum* (Moran *et al.*, 2005).

**Fig 3.**
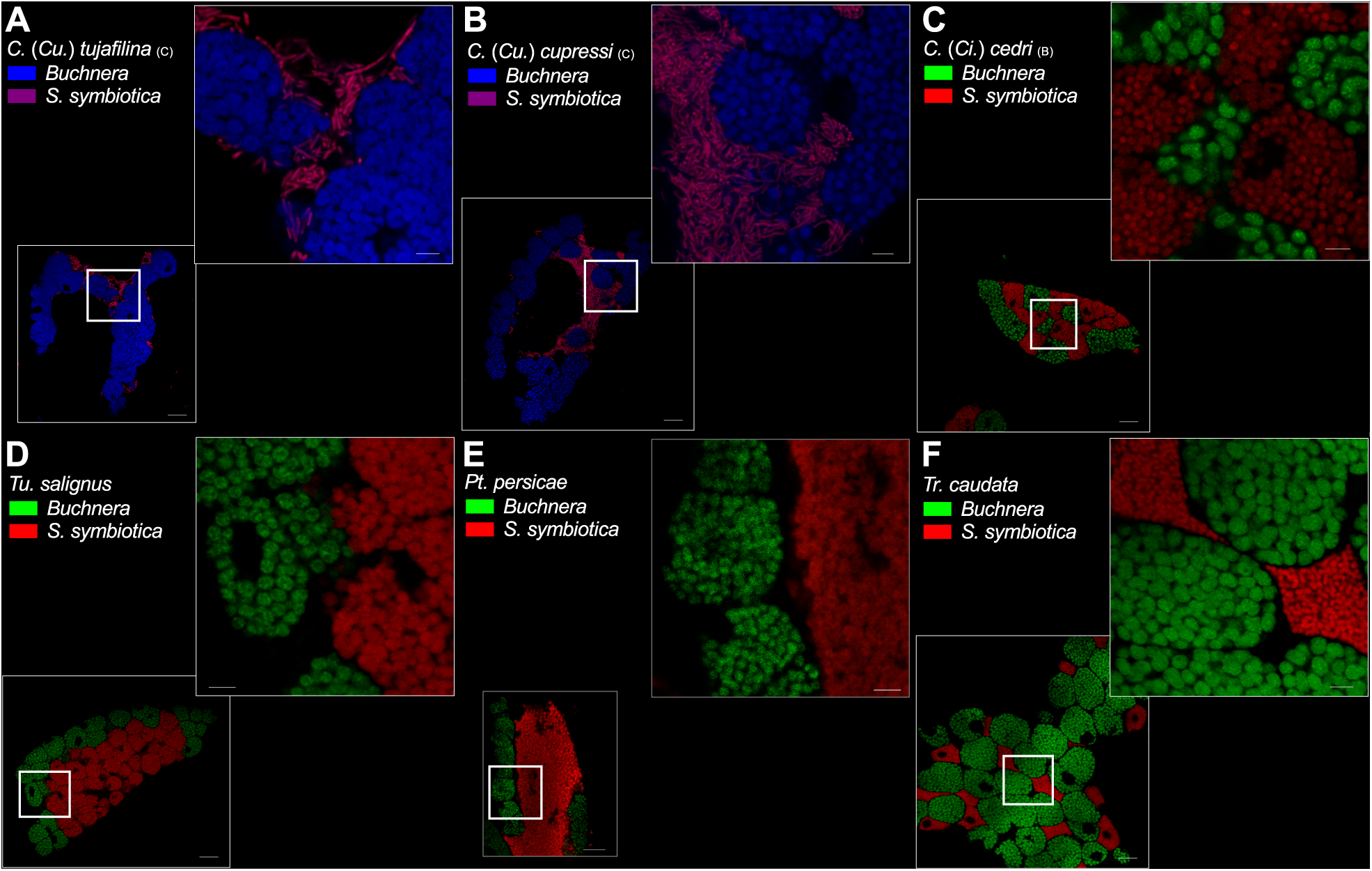
Location and morphology of *S. symbiotica* in selected Lachninae aphids. FISH microscopic images of aphid embryos from selected Lachninae aphids. Symbiont-specific probes were used for FISH, except for panels **A** and **B** in which *Buchnera* was visualised by a general bacterial probe (blue) and *Serratia* by overlapping signals with a *Serratia*-specific probe (red). **(A)** Ventral view of a *C.* (*Cu.*) *tujafilina* bacteriome. **(B)** Lateral-ventral view of a *C.* (*Cu.*) *cupressi* bacteriome. **(C)** Lateral view of a *C.* (*Ci.*) *cedri* bacteriome. **(D)** Lateral view of a *Tu. salignus* bacteriome. **(E)** Lateral-ventral view of a *Pt. persicae* bacteriome. **(F)** Ventral view of a *Tr. caudata* bacteriome. Thick white boxes indicate the magnified region, depicted in the top-right of each panel. The scientific name for each species along with the false colour code for each fluorescent probe and its target group are shown at the top-left of each panel. Scale bars from the unmagnified and magnified FISH images represent 20 and 5µm, respectively.

### *Sodalis*-like secondary symbionts

*Sodalis*-like 16S rRNA gene sequences have been previously amplified from some Lachninae aphids, including *Eulachnus* spp., *Nippolachnus piri*, and *Cinara* (*Cinara*) *glabra* (*Cinara* cluster C) (Burke *et al.*, 2009). Here, we have confirmed, by 16S rRNA gene sequencing, the presence of a ***S****odalis*-**l**ike **s**econdary **s**ymbiont (hereafter **SLSS**) in different populations of *Eulachnus mediterraneus* (Spain) and *Eulachnus rileyi* (Austria, France, and Spain) (**Table S1**). Additionally, using a specific primer designed to target the 16S rRNA gene of the SLSS of *Eulachnus* spp. (see **Materials and Methods**), we detected and sequenced SLSS 16S rRNA gene amplification products from all the collected *Cinara* (*Schizolachnus*) *obscurus* (clade B) populations from Austria, France, and Spain, which were almost identical to each other, pointing towards this symbiont being fixed in this aphid species. Although we lacked enough specimens to perform FISH microscopy, we were able to amplify a sequence from a SLSS from a population of the closely related aphid species *Cinara* (*Schizolachnus*) *pineti*. In this species, we were able to corroborate that the SLSS was the sole secondary bacterium lineage found through a MiSeq amplicon sequencing of the V3-V4 region of the 16S (**Figure 4F**). A similar approach has been recently been used to successfully detect symbionts associated to *Cinara* species (Jousselin *et. al.*, 2016). By means of a Bayesian phylogenetic reconstruction, we determined that SLSSs of Lachninae aphids constitute at least four different lineages, nested within an unresolved clade made up of *Sodalis* bacteria, and *Sodalis*-like symbionts from different insect species (**Fig 4A**). Interestingly, the SLSS from *Eulachnus* spp. form a well-supported monophyletic clade, reinforcing previous results (Burke *et al.*, 2009) and pointing towards a common origin. Considering the close phylogenetic relationship of *Eulachnus* and *Essigella* (**Fig 1**; Chen, Favret, *et al.*, 2015), and the fact that *Essigella* has not been found associated to neither *S. symbiotica*, *Candidatus* Hamiltonella defensa, nor *Candidatus* Regiella insecticola endosymbionts (Russell *et al.*, 2003), we hypothesised that the SLSS detected in *Eulachnus* spp. could have been either fixed in the common ancestor of these two genera or right before the diversification of *Eulachnus* species. Regretfully, we were unable to recover any secondary symbiont’s 16S rRNA gene sequence from *Essigella californica* (collected in France), neither by specific PCR nor by molecular cloning (50 colonies analysed). In the case of the SLSSs from both analysed *Cinara* (*Schizolachnus*) species, we found they were also recovered as a well-supported monophyletic clade, providing evidence towards their common origin.

Given our failure to detect sequence belonging to a secondary symbiont in *Es. californica* using the aforementioned methods, we amplified the V3-V4 region of the 16S rRNA gene and performed massive sequencing in the MiSeq Illumina platform. Surprisingly, we were unable to detect an additional bacterial lineage in *Es. californica* (**Figure S2**). This could reflect either the very low quantity of DNA belonging to the secondary endosymbiotic bacteria (relative to *Buchnera*’s), or a strong bias of the “universal” PCR primers used for this protocol towards amplifying *Buchnera*’s 16S rRNA gene.

**Fig 4.**
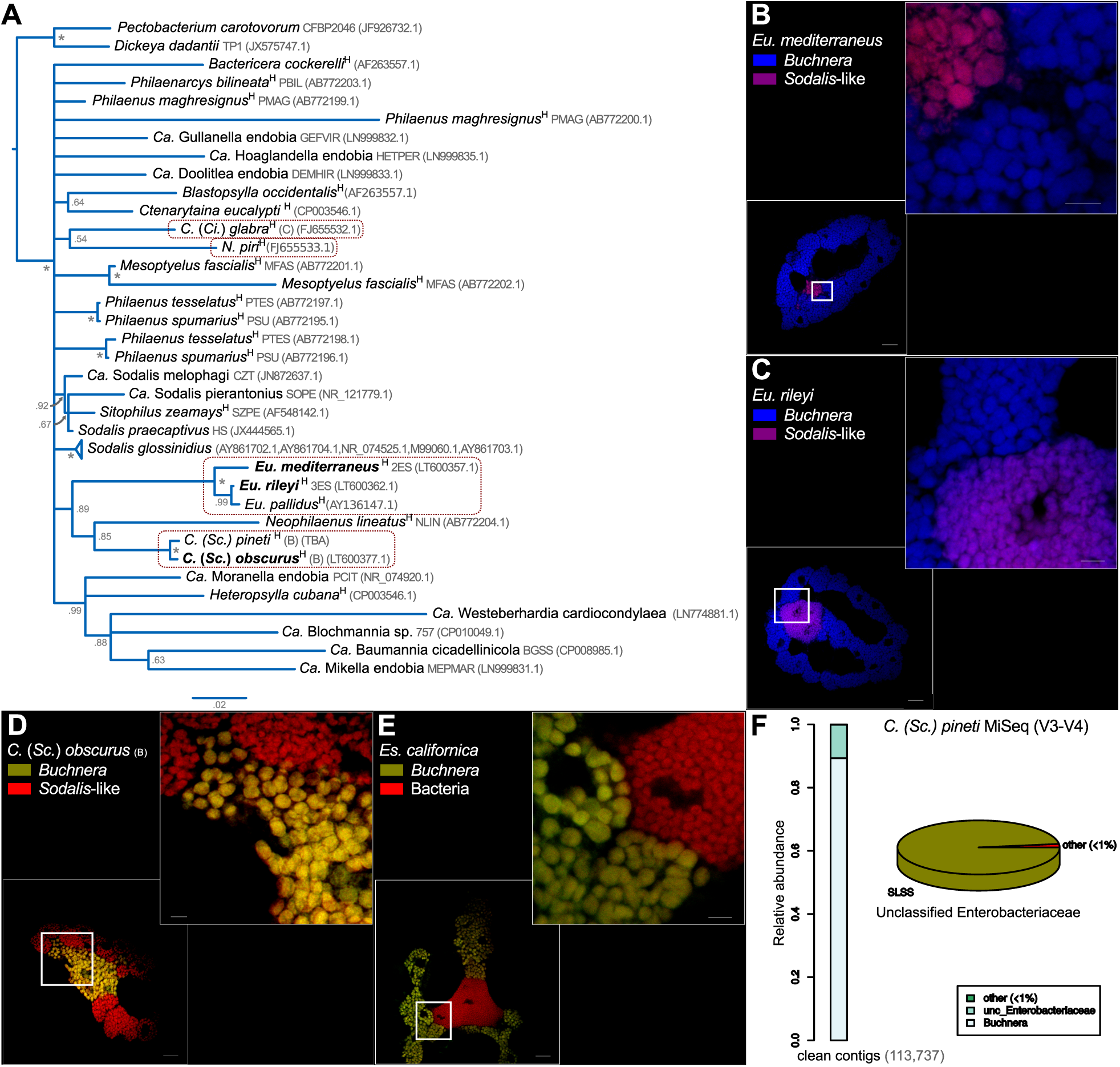
Location and 16S rRNA phylogenetic relationships of SLSS of Lachninae aphids. **(A)** Bayesian phylogram depicting the relationships and placement of known SLSS from aphids. The superscript H at the end of the full species name indicates the symbiont’s host name was used. Strain/isolate names are indicated in grey following the species name. Bold-lettered species names indicate the species selected for FISH microscopy. Values at nodes indicate the posterior probability. An “*” at the node indicate a posterior probability of 1. **(B-E)** FISH microscopic images of aphid embryos of selected Lachninae aphids. In panels **B** and **C**, *Buchnera* was visualised by a general bacterial probe (blue) and SLSSs by overlapping signals with a SLSS-specific probe (red). In panels **D** and **E**, the SLSSs were visualised by a general bacterial probe (red) and *Buchnera* by overlapping signals with a *Buchnera*-specific probe (green). **(B)** Lateral view of an *Eu. mediterraneus* bacteriome. **(C)** Ventral view of an *Eu. rileyi* bacteriome. **(D)** Lateral-ventral view of an *Es. californica* bacteriome. **(E)** Lateral view of a *C.* (*Sc.*) *obscurus* bacteriome. Thick white boxes indicate the magnified region, depicted in the top-right of each panel. The scientific name for each species along with the false colour code for each fluorescent probe and its target group is indicated at the top-left of each panel. Scale bars from the unmagnified and magnified FISH images represent 20 and 5 µm, respectively. **(F)** Stacked bar plot showing the relative abundance of contigs assigned to taxonomical units. On the right, pie chart showing the **MegaBLAST** assignment of V3-V4 contigs to reference 16S rRNA genes from aphid endosymbionts. “<1%” indicates the percentage relative to all the reads (N).

To localise the SLSS within the bacteriome, we used the *Eulachnus* SLSS specific reverse PCR primer as a probe for FISH on dissected aphid embryos (**Table S2**). We found that all individuals from both *Eu. rileyi* and *Eu. mediterraneus* harbour the SLSS inside specific bacteriocytes within the bacteriome (**Figs 4B-C and S1M-O**). Additionally, since we were unable to determine the 16S rRNA gene sequence of the putative secondary endosymbiont of *Es. californica*, we used a combination of a general bacterial and a *Buchnera*-specific probe for FISH in embryos of this aphid species. We observed that there was indeed a distinct secondary bacterial symbiont with a very similar morphology and location as that of *Eulachnus* spp. (**Figs 4D and S1P-R**). In both *Eulachnus* spp. and *Es. californica*, we found that the symbiont is somewhat underrepresented compared to *Buchnera*, similarly to what is observed for the SLSS of *N. piri* (Fukatsu *et al.*, 1998). This could be the reason why we failed to detect it in *Es. californica*, even whith V3-V4 MiSeq amplicon sequencing. Regarding *C.* (*Sc.*) *obscurus*, we did not observe a staining when the SLSS probe (designed for the SLSS of *Eulachnus* spp.) was used for FISH in this aphid species. Therefore, we used the same approach as for *Es. californica*. Using a general bacterial probe in combination with a *Buchnera-* specific probe, we found that *C.* (*Sc.*) *obscurus* harbours two phylogenetically distinct spherical endosymbionts in separate bacteriocytes (**Figs 4E and S1S**). In contrast to *Eulachnus*, the SLSS bacteriocytes from *C.* (*Sc.*) *obscurus* are more abundant and located along the bacteriome surrounding *Buchnera* bacteriocytes.

### “X-type” secondary symbionts

For the current study, we were able to collect two populations of *Ma. submacula* aphids. Since the an “**X-type**” (or **PAXS**) symbiont was suspected to be the secondary symbiont of this aphid species (Lamelas *et al.*, 2008; Burke *et al.*, 2009), we used a specific PCR assay to confirm the presence of this endosymbiont in both populations. Through this assay, we recovered 16S rRNA gene fragments sharing 100% sequence identity with each other and >99% with other X-type symbionts. To facilitate phylogenetic analysis, we additionally performed molecular cloning of the 16S rRNA using universal primers (**Table S2**). Additionally, through the same method, we found an “X-type” symbiont associated with the aphid *C.* (*Ci.*) *confinis*. A Bayesian phylogenetic analysis of the different Aphididae “X-type” symbionts revealed that these form a well-supported monophyletic cluster closely related to *Candidatus* Hamiltonella defensa and *Candidatus* Regiella insecticola, facultative endosymbionts from *Ac. pisum* (**Fig 5A**). Particularly, the sequences obtained from *Ma. submacula* populations from three different countries form a well-supported monophyletic clade (separate from that of *C.* [*Cu.*] *confinis*, *C.* [*Cupressobium*] *juniperi* [clade C], and *Ac. pisum*), and show a high sequence identity among each other (>99%). We then performed FISH analysis on *Ma. submacula* embryos using specific probes for *Buchnera* and X-type (see **Materials and Methods**). We found that all analysed individuals from both *Ma. submacula* populations contained X-type symbionts distributed along the bacteriome, both surrounding *Buchnera* bacteriocytes and in their own distinct ones (**Figs 5B and S1T-U**). Regarding *C.* (*Cu.*) *confinis*, we lacked enough individuals to perform FISH analysis, and therefore its localisation within the bacteriome remains undetermined.

### “*Candidatus* Fukatsuia” gen. nov. and “ *Candidatus* Fukatsuia symbiotica” sp. nov

Given that X-type symbionts form a well-supported monophyletic clade with high sequence identity (>99%), we propose the specific name “***Candidatus* Fukatsuia symbiotica**” for the lineage of enterobacterial symbionts found, so far, affiliated only to aphids (Hemiptera: Aphididae). *Ca.* Fukatsuia’s closest relative, by 16S rRNA gene sequence identity, would be *Budvicia diplopodorum* strain D9 (INSDC accession number HE574451.1), with which it shares 93% sequence identity. The generic name “Fukatsuia” is in honour of Dr. Takema Fukatsu (Prime Senior Researcher at the National Institute of Advanced Industrial Science and Technology, Japan), who has enormously contributed to the study of aphid biology and that of their endosymbionts, with particular emphasis on his early work on secondary endosymbionts form Lachninae aphids. The specific epithet ‘symbiotica’ alludes to the symbiotic habit of *Ca.* Fukatsuia bacteria.

In the Lachninae aphid *Ma. submacula*, “*Ca.* Fukatsuia” is found inhabiting the bacteriome tissue and its cell presents a filamentous shape of variable length. Its tissue localisation and cell shape in aphids other than *Ma. submacula* remains unknown. Similarly to *S. symbiotica*, different *Ca.* Fukatsuia lineages are of different dispensability to their hosts: being a defensive facultative symbiont (of variable degrees of protection, depending on the strain) in *Ac. pisum* (Guay *et al.*, 2009; Heyworth and Ferrari, 2015), and being a putative co-obligate symbiont in *Ma. submacula* aphids. Co-obligate lineages of “*Ca.* Fukatsuia symbiotica” may well represent separately evolving units, and thus, some lineages may constitute separate species within the same genus. This could be the case for the well-supported specific lineage associated to *Ma. submacula* aphids, however further genome data from several “*Ca.* Fukatsuia” symbionts is needed to test this hypothesis. Currently available sequences that correspond to “*Ca.* Fukatsuia symbiotica” are deposited under INSDC accession numbers FJ821502.1, KP866544.1, KP866545.1, LT600381.1, EU348311.1, EU348312.1, FJ655539.1, LT600338.1, and LT600340.1.

**Fig 5.**
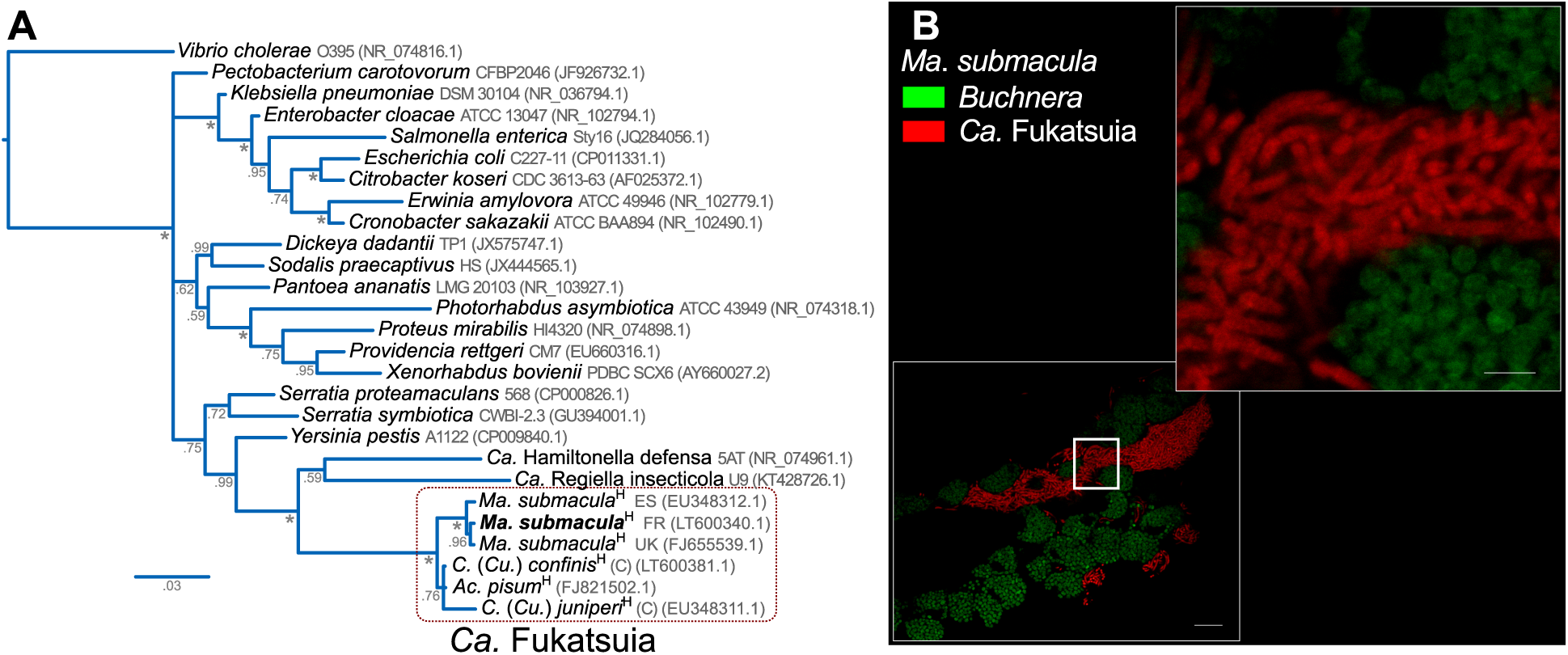
16S rRNA based phylogenetic relationships of *Ca.* Fukatsuia and location of *Ca.* Fukatsuia symbiotica in *Ma. submacula*. **(A)** Bayesian phylogram depicting the relationships and placement of the currently available *Ca.* Fukatsuia from aphids and selected Enterobacteriaceae, using *Vibrio cholerae* as an outgroup. The superscript H indicates that the symbiont’s host name was used. **(B)** FISH microscopic images of a lateral view of a *Ma. submacula* bacteriome. Thick white box indicates the magnified region, depicted in the top-right of the panel. The scientific name for the species along with the false colour code for each fluorescent probe and its target group is shown at the top-left of the panel. Scale bars from the unmagnified and magnified FISH images represent 20 and 5 µm, respectively.

## Discussion

Many insects maintain intimate associations with obligate endosymbiotic bacteria harboured in specialised organs (bacteriomes). One crucial question within the field is how do these associations evolve. One way of approaching this question is through the study of “recently” acquired endosymbionts. In the Lachninae subfamily of aphids, it has previously been proposed that an ancient loss of the riboflavin biosynthetic capability by *Buchnera* promoted the settlement of a co-obligate secondary endosymbiont (putatively *S. symbiotica*) in the LLCA (Manzano-Marín and Latorre, 2014; Manzano-Marín *et al.*, 2016). Yet, various extant members of this subfamily have been found to be associated to bacterial taxa phylogenetically distinct from *S. symbiotica* (Lamelas *et al.*, 2008; Burke *et al.*, 2009). Therefore, the study of secondary endosymbionts within this subfamily is expected to give important clues regarding symbiont establishment and transition from facultative to co-obligate relationships. However, the studies on the symbiotic systems of aphids belonging to the Lachninae have been hampered by the lack of an evolutionary framework to correctly interpret the genomic and metabolic changes, as well as their links with the different stages of the transformation process towards an obligate intracellular lifestyle (Pérez-Brocal *et al.*, 2006; Lamelas, Gosalbes, Manzano-Marín, *et al.*, 2011; Lamelas, Gosalbes, Moya, *et al.*, 2011; Manzano-Marín and Latorre, 2014).

In this work, we have explored the diversity, phylogenetic relationships, and location of different secondary endosymbionts within key members of the Lachninae subfamily. This has enabled us to propose an evolutionary scenario for the settlement, “stable” internalisation into distinct bacteriocytes, and replacements of the original secondary co-obligate endosymbiont from the LLCA (**Fig 6**). Firstly, with a combination of specific PCR assays, 16S rRNA gene sequencing, and FISH microscopy, we determined that all analysed specimens indeed harbour fixed secondary endosymbionts at the population/species level. This fact, in combination with previously published microscopic (Buchner, 1953; Fukatsu and Ishikawa, 1998; Fukatsu *et al.*, 1998; Lamelas *et al.*, 2008; Pyka-Fościak and Szklarzewicz, 2008; Michalik *et al.*, 2014) and molecular (Pérez-Brocal *et al.*, 2006; Burke *et al.*, 2009; Lamelas, Gosalbes, Manzano-Marín, *et al.*, 2011; Lamelas, Gosalbes, Moya, *et al.*, 2011; Manzano-Marín and Latorre, 2014; Manzano-Marín *et al.*, 2016) data from Lachninae aphids, provides strong evidence for the dependence of members of this subfamily on co-obligatory secondary endosymbionts, putatively due to the ancient pseudogenisation of the riboflavin biosynthetic genes in the *Buchnera* harboured by the LLCA, which lived at least some 85-106 Mya. The detection of *S. symbiotica* in different aphid species from at least six Lachninae genera across all five tribes, along with the genomic data from three strains of this symbiont at different stages of the genome reduction process (Lamelas, Gosalbes, Manzano-Marín, *et al.*, 2011; Manzano-Marín and Latorre, 2014; Manzano-Marín *et al.*, 2016), point towards an early establishment of *S. symbiotica* as co-obligate in the LLCA. While spherical and found consistently inside bacteriocytes (with a highly reduced genome) in *Tu. salignus* and *C.* (*Ci.*) *cedri*, the filamentous and broadly distributed (with a mildly-reduced genome) *S. symbiotica* from *C.* (*Cu.*) *tujafilina* would preserve the traits of the putative “ancient” *S. symbiotica* from the LLCA. This hypothesis would be consistent with the high level of genomic, metabolic, and phenotypic similarity of the co-obligate *S. symbiotica* from *C.* (*Cu.*) *tujafilina* and the facultative *S. symbiotica* from *Ac. pisum* (Moran *et al.*, 2005; Lamelas *et al.*, 2008; Burke and Moran, 2011; Manzano-Marín and Latorre, 2014). We find this scenario to be most parsimonious, as it would require one single event of infection with a shared *S. symbiotica* ancestor in the LLCA followed by at least four “stable” internalisations of *S. symbiotica* into bacteriocytes. This ancient secondary symbiont would have then undergone at least six independent events of symbiont replacement. An alternative scenario would require additional events of symbiont replacement with distinct *S. symbiotica* strains in specific Lachninae lineages. This last scenario is suggested by the lack of general congruency between *Buchnera* and *S. symbiotica* 16S rRNA gene phylogenies and by the multiple *S. symbiotica* lineages recovered from the 16S rRNA gene phylogeny in **Fig 2B**. However, this pattern also suggests different rates of sequence evolution of the *S. symbiotica* symbionts, possibly driven by changes in the *Buchnera*-*S. symbiotica* relationship, such as metabolic pathway splits (e.g. tryptophan) and/or changes in the symbiont’s tissue tropism (Manzano-Marín *et. al.*, 2016). We expect that further sequencing of complete genomes from several Lachninae aphids will help clarify this.

**Fig 6.**
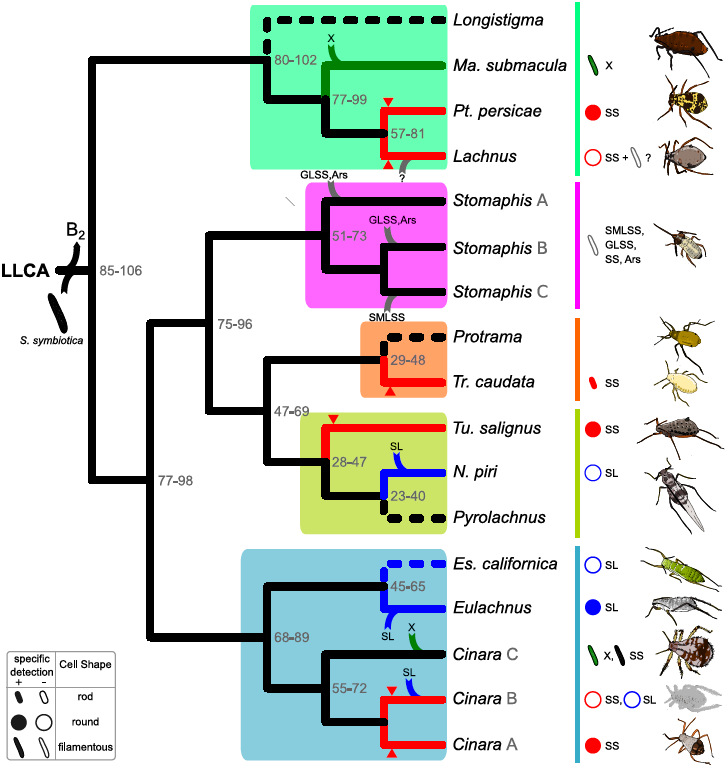
Proposed evolutionary scenario for the establishment, “stable” internalisation, and replacement of secondary co-obligate endosymbionts across the Lachninae. Cladogram displaying the relationships of Lachninae lineages by genera. Coloured boxes shading monophyletic clades as well as vertical lines on the right side delimit the five tribal clades (as depicted in **Fig 1**). Divergence time range estimates (in Mya, and showed at tree nodes) are based on (Chen, Favret, *et al.*, 2015). Incoming lines on branches symbolise the acquisition/replacement of co-obligate secondary symbionts. The outgoing line at the root of the tree stands for the loss of the riboflavin (B2) biosynthetic genes in the *Buchnera* from the LLCA. Green, blue, and grey branches represent lineages where *Ca.* Fukatsuia, a SLSS, or other bacterial symbiont have replaced the original *S. symbiotica* symbiont, respectively. Red branches with an arrowhead pointing to them reflect the “stable” internalisation of *S. symbiotica* into distinct bacteriocytes. At the leaves, shapes symbolising the bacterial endosymbionts’ cell shapes according to the key (bottom-left) and cartoons of selected aphids form the different Lachninae genera are showed. SS= *S. symbiotica*, X=*Ca.* Fukatsuia, Ars=*Arsenophonus*, SMLSS=*Serratia marcescens*-like secondary symbiont, GLSS=*Gilliamella*-like secondary symbiont, SL= SLSS, ?=unknown.

Within the Lachnini tribe, there could have been either one or two independent events of “stable” internalisation and confinement of *S. symbiotica* into distinct bacteriocytes. The latter hypothesis is supported by the lack of congruency between *Buchnera* and *S. symbiotica* lineages from *Pt. persicae* and *Lachnus* spp., suggesting separate events of genome reduction of *S. symbiotica* in these two aphid lineages. Regarding the *Lachnus* genus, microscopic observations of bacteriocytes from *La. roboris* have revealed that it keeps a vertically transmitted association with two spherical-shaped bacteria (presumably *Buchnera* and *S. symbiotica*) residing in separate bacteriocytes and a third filamentous bacterial symbiont residing in distinct bacteriocytes, whose identity remains unknown (Buchner, 1953). If this tertiary symbiotic bacterium was established in the common ancestor of *La. roboris* and *La. quercihabitans*, it could explain the longer branches, relative to *La. tropicalis* (**Fig 2A** and **B**), given that the presence of an additional symbiont could facilitate the process of genomic erosion in the *S. symbiotica* symbiont. Such a phenomenon was observed in those *Buchnera* strains from Lachninae species that have established co-obligate associations with *S. symbiotica* (Manzano-Marín *et al.*, 2016).

Also, we have confirmed that *Ma. submacula* aphids indeed harbour *Ca.* Fukatsuia bacteria (which belong to the group of symbionts previously referred to as “X-type” or PAXS) and determined its location within aphid embryos. We found that *Ca.* Fukatsuia was present in all of the microscopy analysed individuals, which, in combination with previous analyses detecting the presence of these symbionts in a Spanish (Lamelas *et al.*, 2008), and a UK (Burke *et al.*, 2009) population, points towards its obligate status. The morphology and location of *Ca.* Fukatsuia (**Fig 5B and S1T-U**) resembles that observed for facultative endosymbionts of other aphids (Moran *et al.*, 2005), similarly to what is observed for the co-obligate *S. symbiotica* from *C.* (*Cu.*) *tujafilina* (Lamelas *et al.*, 2008; Manzano-Marín and Latorre, 2014). This suggests that *Ca.* Fukatsuia from *Ma. submacula* has not yet undergone a massive genome reduction, contrary to what is observed in the pleomorphic *S. symbiotica* of *C.* (*Ci.*) *cedri* (Lamelas, Gosalbes, Manzano-Marín, *et al.*, 2011) and *Tu. salignus* (Manzano-Marín *et al.*, 2016). This, in combination with the lack of a *S. symbiotica* endosymbiont, points toward a replacement of the “ancient” secondary co-obligate endosymbiont which occurred at least some 77-99 Mya in the branch leading to *Ma. submacula*. It is important to note that *Buchnera* strains from aphids identified as *Ma. submacula* form at least two phylogenetically distinct lineages (**Fig 2A**): one sister to *Maculolachnus sijpkensis* and one sister to this *Ma. submacula*+*Ma. sijpkensis* clade. Thus, in the current work, we refer to the latter as the one associated to *Ca.* Fukatsuia. Given that no *cox1* gene sequence is provided for the *Ma. submacula* whose *Buchnera* strain is recovered as sister to that of *Ma. sijpkensis*, we are unable to judge if these the two *Buchnera* lineages have been indeed isolated form the same species or if the polyphyly of *Buchnera* strains from *Ma. submacula* is due to wrong taxonomic identification.

Regarding the Stomaphidini, we postulate that at least two events of symbiont replacement have occurred. *Stomaphis pini* and *Stomaphis quercisucta* (both belonging to the same clade: *Stomaphis* clade A) have been found to be associated with *S. symbiotica* (Burke *et al.*, 2009; Chen, Wang, *et al.*, 2015), and microscopic investigations into *St. quercus* have revealed that this species houses three vertically-transmitted endosymbiotic bacteria: *Buchnera* plus two secondary symbionts which apparently reside inside the same secondary bacteriocytes (Buchner, 1953; Pyka-Fościak and Szklarzewicz, 2008). These could be an *Arsenophonus* and/or a *Gilliamella*-like secondary symbiont (**Figure S3**), both of which have been found associated to different Polish populations of this aphid species (Burke *et al.*, 2009). In addition, both *Stomaphis aphananthae* and *St. yanonis* are associated with SMLSSs (Burke *et al.*, 2009), suggesting an establishment of this symbiont in the branch leading to the clade comprising these two species (the unresolved *Stomaphis* clade C). Furthermore, microscopic analyses of *St. yanonis* bacteriomes have revealed this species indeed houses a tubular secondary symbiont, putatively SMLSS, in separate bacteriocytes located on the surface of the bacteriome “core” formed by the *Buchnera* bacteriocytes (Fukatsu and Ishikawa, 1993, 1998). Consequently, we propose at least two events of acquisition of a new endosymbiont: one before the diversification of the clade comprising *St. cupressi* (*Stomaphis* clade A) and another one before the expansion of the large unresolved clade including *St. yanonis* (Stomaphis clade C) (**Fig 2A**).

With respect to the Tramini+Tuberolachnini clade, all currently analysed members have been found to be associated with *S. symbiotica* (Burke *et al.*, 2009), except for *N. piri*, which contains a putatively pleomorphic SLSS housed inside separate bacteriocytes (Fukatsu and Ishikawa, 1993, 1998; Fukatsu *et al.*, 1998; Burke *et al.*, 2009). In the case of both *Tu. salignus* and *Tr. caudata* (which started diverging at least some 47-69 Mya), the *S. symbiotica* symbiont is found exclusively within bacteriocytes (**Figs 3D, F, S1H-I,** and **K-L**). However, the cell shape of the endosymbionts is strikingly different. While in *Tu. salignus* they occur as large spherical-shaped cells, in *Tr. caudata* the symbionts have a small rod-shaped morphology, resembling the cell shape of free-living *Serratia* strains. This could be indicative of this bacterium being in the very first stages of “stable” internalisation into bacteriocytes, while still preserving its rod shape and putatively having a genome resembling closely that of *S. symbiotica* from *C.* (*Cu.*) *tujafilina*, rather than that of *Tu. salignus*.

In regards to the Eulachnini, most *Cinara* spp. have been consistently found associated to *S. symbiotica* strains. Microscopic investigations of *Cinara* (*Cinara*) *pini* (clade A) and *C.* (*Ci.*) *cedri* have revealed they indeed harbour a pleomorphic secondary endosymbiotic bacterium obligatorily inside bacteriocytes (Fukatsu *et al.*, 1998) (**Figs 3C and S1E-G**), and in the case of the latter, genomic-based metabolic inference has established that both *Buchnera* and *S. symbiotica* are required for the biosynthesis of various essential nutrients (Gosalbes *et al.*, 2008; Lamelas, Gosalbes, Manzano-Marín, *et al.*, 2011). Additionally, a high level of congruency between the phylogenetic relationships of *Buchnera* and *S. symbiotica* strains from clade A *Cinara* (**Fig 2A** and **B**) suggests a single event of drastic genome reduction followed by divergence, similar to what is observed for *Buchnera*. On the contrary, *S. symbiotica* from clade B *Cinara* do not show this congruent pattern, pointing possibly to independent events of drastic genome reduction. Within *Cinara* clade B, *C. (Sc.) obscurus* would represent a case of symbiont replacement by a SLSS, which is present obligatorily inside bacteriocytes (**Fig 4E and S1S**). This SLSS is also present as in the closely related *C.* (*Sc.*) *pineti* (in which it is the only other symbiont present), and according to a previous study, this species also presents a spherical secondary endosymbiotic bacterium (presumably the detected SLSS) which is vertically transmitted in both oviparous and viviparous generations (Michalik *et al.*, 2014). Taken together, this suggests the replacement of *S. symbiotica* by a SLSS in the common ancestor of these two *Cinara* (*Schizolachnus*) species. Whether or not this symbiont is widespread within the *Cinara* (*Schizolachnus*) subgenus remains to be explored. As most *Cinara*, *C.* (*Cu.*) *tujafilina* and *C.* (*Cu.*) *cupressi*, are associated to *S. symbiotica* strains. However, both the location and 16S rRNA gene sequence of these more closely resemble the facultative strains from Aphidinae aphids (**Figs 2B, 3A-B, S1A-D**; Moran *et al.*, 2005). Genome-based metabolic inference has provided evidence towards the obligate status of *S. symbiotica* in *C.* (*Cu.*) *tujafilina*, given the loss of the riboflavin biosynthetic capability of *Buchnera*, an essential co-factor now synthesised by *S. symbiotica* (Manzano-Marín and Latorre, 2014). This, in addition to the consistent association of these two *Cinara* (*Cupressobium*) species with *S. symbiotica*, led us to infer that these aphids do indeed keep an obligate association with closely related secondary endosymbiotic strains. Within *Cinara* clade C, evidence of at least one species being affiliated to a SLSS (**Fig 4A**, *Cinara* [*Ci.*] *glabra*) and two to *Ca.* Fukatsuia (**Fig 5A,** *Cinara* [*Cu.*] *confinis and Cinara*[*Cu.*]*juniperi*), rather than *S. symbiotica*, suggests that some events of symbiont replacement have occurred in this group of species. This could have been facilitated due to the niche occupied by *S. symbiotica*, being similar to that of facultative endosymbionts of *Ac. pisum* (Sandström *et al.*, 2001; Moran *et al.*, 2005; Sakurai *et al.*, 2005; Tsuchida *et al.*, 2005, 2010). Finally, we propose a symbiont replacement event by a SLSS in the branch leading to the *Eulachnus* species. Consistent with previous observations in *Eu. rileyi* (Michalik *et al.*, 2014), we found that both *Eu. rileyi and Eu. mediterraneus* species harbour spherical SLSSs in separate bacteriocytes, spatially-arranged in a very similar fashion (**Fig 4B, C, S1M, O**). Also, even though we were unable to recover a 16S rRNA gene sequence belonging to a bacterial taxon other than *Buchnera* in *Es. californica* (in a greater abundance than 1%), we were able to detect by FISH microscopy the presence of spherical bacterial endosymbionts residing in distinct bacteriocytes, localised similarly to those inhabited by SLSSs in *Eulachnus* spp. (**Fig 4D, SR**). Therefore, pending further studies, it could be suggested that the secondary symbiont found in *Es. californica* could belong to the same lineage as the SLSSs of *Eulachnus* species.

A feature we observed repeatedly was the change in the endosymbionts’ tissue tropism along distantly related aphid species. This changes included both bacteriocyte arrangement within the bacteriome and “stable” internalisation of the secondary endosymbionts in distinct bacteriocytes. In the case of *S. symbiotica* and *Buchnera*, genome data is available for three species. Regarding the bacteriocyte arrangement within the bacteriome, a similar case has been previously reported within spittlebugs (Auchenorrhyncha: Cercopoidea). Within this superfamily, species belonging to the tribe Philaenini have undergone symbiont replacement of the ancient *Candidatus* Zinderia endosymbiont by a *Sodalis*-like symbiont (Koga *et. al.*, 2013). This shift in co-obligate symbiont is also accompanied by a newly evolved type of bacteriocyte with a different arrangement. In Lachninae aphids, changes in the bacteriocyte arrangement within the bacteriome could also be linked to symbiont replacement events, and/or changes in the interdependent metabolic “wiring” of their symbionts which promotes “stable” internalisation into distinct bacteriocytes in a non-deterministic matter. Regarding the latter, we have previously postulated that these changes in the interdependent metabolic “wiring” could also be involved in the “stable” internalisation into distinct bacteriocytes (Manzano-Marín *et. al.*, 2016), which speculatively could be triggered by the constraints on the exchange of certain intermediary metabolites. Developmentally, this shift in tissue tropism would involve a change in the development and colonisation of the bacteriome by the secondary symbionts. In *Ac. pisum*, *S. symbiotica* acquires a broad distribution in the bacteriome (e.g. occupying both sheath cells, secondary bacteriocytes, and co-infecting *Buchnera*’s) following the formation of sheath cells (Koga *et. al.*, 2012). Before this, and after bacteriocyte cellularisation and symbiont sorting, the two “stably” internalised symbionts of the different Lachninae aphids would putatively remain confined to their own bacteriocytes until vertical symbiont transmission.

In summary, we propose an evolutionary framework which should assist in future studies on the Lachninae, a symbiont-diverse subfamily. Our findings reveal a dynamic pattern for the evolutionary history of “recently” established endosymbionts, thus contributing to a better understanding of how mutualism in endosymbiotic associations can evolve. We believe that further studies directed towards the bacteriome development and its colonisation by endosymbiotic bacteria in species from the Lachninae subfamily, could also provide hints towards the evolution of new spatial arrangements and even type of bacteriocytes. The role these recently-acquired bacteria have played in the adaptation of their aphid hosts to different niches/feeding sites/plants and their role in speciation in this peculiar subfamily remains to be explored.

## Materials and Methods

### Aphid collection and storage

All aphids used for this study were collected in various locations around Rennes (France), Vienna (Austria), and Valencia (Spain). Collection details can be found in **Table S1**. Aphids used for DNA extraction were stored at-20°C in absolute ethanol inside a 1.5 mL Eppendorf tube. Aphids used for fluorescence *in situ* hybridisation experiments were dissected in absolute ethanol to extract embryos. These were then directly transferred to modified Carnoy’s fixative (6 chloroform : 3 absolute ethanol : 1 glacial acetic acid) and left overnight, following (Koga *et al.*, 2009) protocol to quench autofluorescence. Briefly, fixed embryos were washed with absolute ethanol and transferred into a 6% solution of H2O2 diluted in absolute ethanol and were then left in this solution for two to six weeks (changing the solution every three days). When bleached, they were washed twice with absolute ethanol and stored at-20°C.

### Fluorescence *in situ* hybridisation

Hybridisation of aphid embryos was performed overnight at 28°C in standard hybridisation buffer (20mM Tris-HCl [pH 8.0], 0.9 M NaCl, 0.01% SDS, and 30% formamide) and then washed (20mM Tris-HCl [pH 8.0], 5mM EDTA, 0.1 M NaCl, and 0.01% SDS) before slide preparation. The slides were examined using a confocal laser scanning microscope (TCS SP5 X, Leica; and FV1000, Olympus). A list of specific probes used for each aphid species is available in **Table S2**. *Buchnera*, *S. symbiotica*, and *Ca.* Fukatsuia competitive probes were designed based on (Gómez-Valero *et al.*, 2004) and adapted to match the target bacterial strain. The *Eulachnus* SLSS probe was designed based on (Attardo *et al.*, 2008), adapted to match the target strain. The embryos from at least 10 individuals were analysed per sample.

### 16S rRNA gene PCR, cloning, and sequencing

Since all endosymbionts detected in Lachninae members so far are bacteria, we used the primers 16SA1 and 16SB1 (Fukatsu and Nikoh, 1998) to amplify partial 16S rRNA genes (*circa* 1.5 kbp) for cloning and sequencing. This strategy was adopted in selected cases to facilitate phylogenetic reconstruction. Resulting amplicons were cloned into the pGEM-T Easy Vector (Promega) and SP6 (5’-ATTTAGGTGACACTATAG-3’) and T7 (5’-TAATACGACTCACTATAGGG-3’) primers were used for amplification and sequencing of the cloned DNA (at least 5 clones from each species). Specific primers for either *Buchnera* or *S. symbiotica* were designed based on the FISH probes. Specific PCR reactions and sequencing were done mainly to confirm the presence of the secondary endosymbionts. In the case of the SLSS and *Ca.* Fukatsuia, specific primers were designed based on (Attardo *et al.*, 2008) (“ *Sodalis* specific”) and (Guay *et al.*, 2009) (“PAXSF”), respectively. For a full list of primers pairs and PCR conditions see **Table S2**. All sequences have been uploaded to the European Nucleotide Archive and are pending accession (temporarily available at https://figshare.com/s/8eb9686e546394547fe4).

### MiSeq sequencing of the V3-V4 region of the 16S rRNA gene from bacteria associated to *Es. Californica* and *C.* (*Sc.*) *pineti*

Using the same DNA extracted for PCR and cloning, amplification and sequencing of the V3-V4 region (using standard Illumina primers 5’-CCTACGGGNGGCWGCAG-3’ and 5’- GACTACHVGGGTATCTAATCC-3’) of the 16S rRNA gene was performed in an Illumina MiSeq machine (paired-end 2x300 bp) at the FISABIO Center (Generalitat Valenciana). Next, **mothur** v1.31.2 (Schloss *et. al.*, 2009) was used for merging of the paired-ends reads and taxonomic assignment of the resulting contigs. Briefly, reads where first quality trimmed using **fastx_toolkit** v0.0.14 (http://hannonlab.cshl.edu/fastx_toolkit/, last accessed November 9, 2016). Next, we joined the overlapping paired ends reads using the *make.contigs* function of **mothur** and filtered out all contigs shorter than 420 bps. Then, contigs were aligned to the **SILVA NR99** v128 database (Quast *et. al.*, 2013) and those with less than 90% of their length aligned were filtered out. After another step of redundancy removal, rare sequences (possibly resulting from sequencing errors) were merged with frequent unique sequences with a mismatch no greater than 2 bp (*precluster* function). Resulting contigs were then screened for chimeric sequences using (*chimera.uchime* function). Remaining contigs were then taxonomically assigned to genus level using the *classify.seqs* function and the **SILVA NR99** v128 database (cutoff=90). Lastly, the remaining unclassified sequences (namely unclassified Enterobacteriaceae) that exceeded 1% of the total contigs were used for a **MegaBLAST**(Zhang *et. al.*, 2004) search against a database of representatives including all 16S aphid endosymbiont sequences available to date. The best hit for each read, if this surpassed 94.5% identity, was used for genus-level assignment.

### Phylogenetic analyses

All phylogenetic analyses were performed as follows. First **SSU-ALIGN** v0.1 (Nawrocki, 2009) was used to align 16S rRNA sequences, followed by visual inspection of the alignments in **AliView** v1.17.1 (Larsson, 2014). Then, **GBlocks** v0.91b (Castresana, 2000) was used to eliminate poorly aligned positions and divergent regions with the option ’-b5=h’ to allow half of the positions with a gap. The final alignments were transformed into nexus format (available online at https://figshare.com/s/8eb9686e546394547fe4) for phylogenetic analysis in **MrBayes** v3.2.5 (Ronquist *et al.*, 2012) under the GTR+I+G model. Two independent runs, each with four chains (three “heated”, one “cold”), were run for 5,000,000 generations discarding the first 25% as burn-in and checked for convergence. Visualisation and tree-editing was done in **FigTree** v1.4.1 (http://tree.bio.ed.ac.uk/software/figtree/, last accessed November 9, 2016) and **Inkscape** v0.91 (http://www.inkscape.org/en/, last accessed November 9, 2016), respectively. For a full list of accession numbers of sequences used for phylogenetic analyses see **Table S3**.

## Acknowledgements

The authors would like to acknowledge emeritus professor José Manuel Michelena Saval, professor Joaquin Baixeras, Nicolás Pérez Hidalgo, Christelle Buchard, and Evelyne Turpeau for their invaluable and expert help in collection and/or taxonomic identification of aphid samples. Also, we thank and acknowledge artist/scientist Jorge Mariano Collantes Alegre for the aphid cartoons in **Fig 6**. This work has been funded by the Ministerio de Economía y Competitividad (Spain) co-financed by FEDER funds [BFU2015-64322-C2-01-R to A.L.]; the European Commission [Marie Curie FP7 PITN-GA-2010-264774-SYMBIOMICS to A.M.M]; the Consejo Nacional de Ciencia y Tecnología (Mexico) [Doctoral scholarship CONACYT 327211/381508 to A.M.M.]; and the Austrian Science Fund [project no. P22533-B17 to M.H.]. The Plant Health and Environment department of INRA is also acknowledged for financial support to JC Simon. The funders had no role in study design, data collection and analysis, decision to publish, or preparation of the manuscript. The authors declare no conflict of interest.

## Supporting Information

**S1 Table. Accession numbers, collection data, and taxonomic status of sampled aphids and their endosymbionts.**

**S2 Table. Primers and probes used to detect endosymbionts from the different aphid species.** Primers and probes used in this study to detect endosymbionts and their specificities.

**S3 Table. Accession numbers of sequences used for phylogenetic reconstruction.**

**S1 Fig. Location and morphology of secondary symbionts in selected Lachninae aphids.** FISH microscopic images of aphid embryos from selected Lachninae aphids. Symbiont-specific probes were used for FISH, except for panels **A-D** and **M-S** in which either one of the symbionts was visualised by a general bacterial probe and the other. Colors are as in corresponding images of the same aphid species in the main text. **(A)** Lateral and **(B-C)** Dorsal views of a *C.* (*Cu.*) *tujafilina* bacteriome of an early and later embryos, respectively. **(D)** Lateral view of a *C.* (*Cu.*) *cupressi* bacteriome of an early embryo. **(E)** Lateral and **(F-G)** Dorsal views of a *C.* (*Ci.*) *cedri* bacteriome of an earlier and later embryos, respectively. **(H)** Lateral and **(I)** ventral view of a *Tu. salignus* bacteriome of an earlier and a late embryo. **(J)** Lateral-ventral view of a *Pt. persicae* bacteriome. Lateral and **(L)** Ventral view of a *Tr. caudata* bacteriome of an early and later embryo. **(M)** Ventral view of an *Eu. mediterraneus* bacteriome. **(N)** Lateral and **(O)** ventral view of an *Eu. rileyi* bacteriome of an early and later embryo. **(P)** Lateral and **(Q-R)** ventral-lateral views of an *Es. californica* bacteriome from an early and later embryos. **(S)** View of an early embryo of a *C.* (*Sc.*) *obscurus* bacteriome. **(T)** Dorso-lateral and **(U)** lateral views of early embryos of a *Ma. submacula* bacteriome. Thick white boxes indicate the magnified region, depicted in the top-right of each panel. The scientific name for each species along with the false colour code for each fluorescent probe and its target group are shown at the top-left of each panel. Scale bars from the unmagnified and magnified FISH images represent 20 and 5µm, respectively.

**S2 Fig. 16S V3-V4 rRNA amplicon sequences taxonomic assignment.** Stacked bar plot showing the relative abundance of contigs assigned to taxonomical units. On the right, pie chart showing the **MegaBLAST** assignment of V3-V4 contigs to reference 16S rRNA genes from aphid endosymbionts. “<1%” indicates the percentage relative to all the reads (N).

**S3 Fig. 16S rRNA gene-based phylogenetic relationships of GLSS strains from the Aphididae.** Bayesian phylogram depicting the relationships and placement of the currently available GLSS strains from Aphididae and selected Enterobacteriaceae, Pasteurellaceae, and Orbaceae. The superscript H at the end of the full species name indicates the symbiont’s host name was used. The accession numbers for each sequence used is indicated within parenthesis after the strain name.

